## Supplemental Figures for "Integrator complex subunit 12 knockout overcomes a transcriptional block to HIV latency reversal"

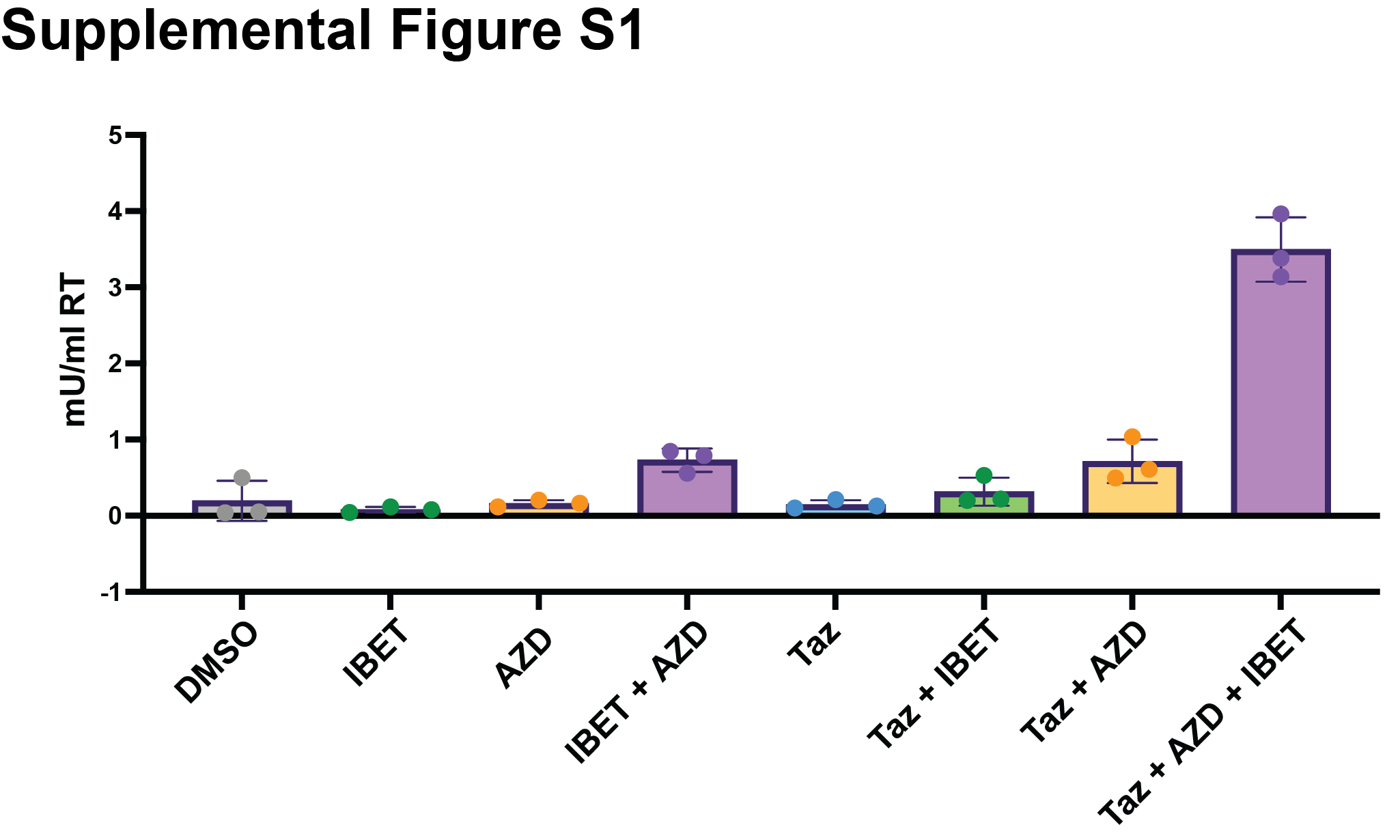

**Figure 1—figure supplement 1**: **Validation of HIV-CRISPR screen with inhibitors.** J-Lat 5A8 cells were knocked out for AAVS1 and then treated with DMSO, 100 nM I-BET151 (IBET) for 48 hrs, 10 nM AZD5582 (AZD) for 48 hrs or 10 uM the EZH2i Tazemetostat (Taz) for 96 hrs. HIV reverse transcriptase activity was measured from the supernatant (reported in mU/mL).

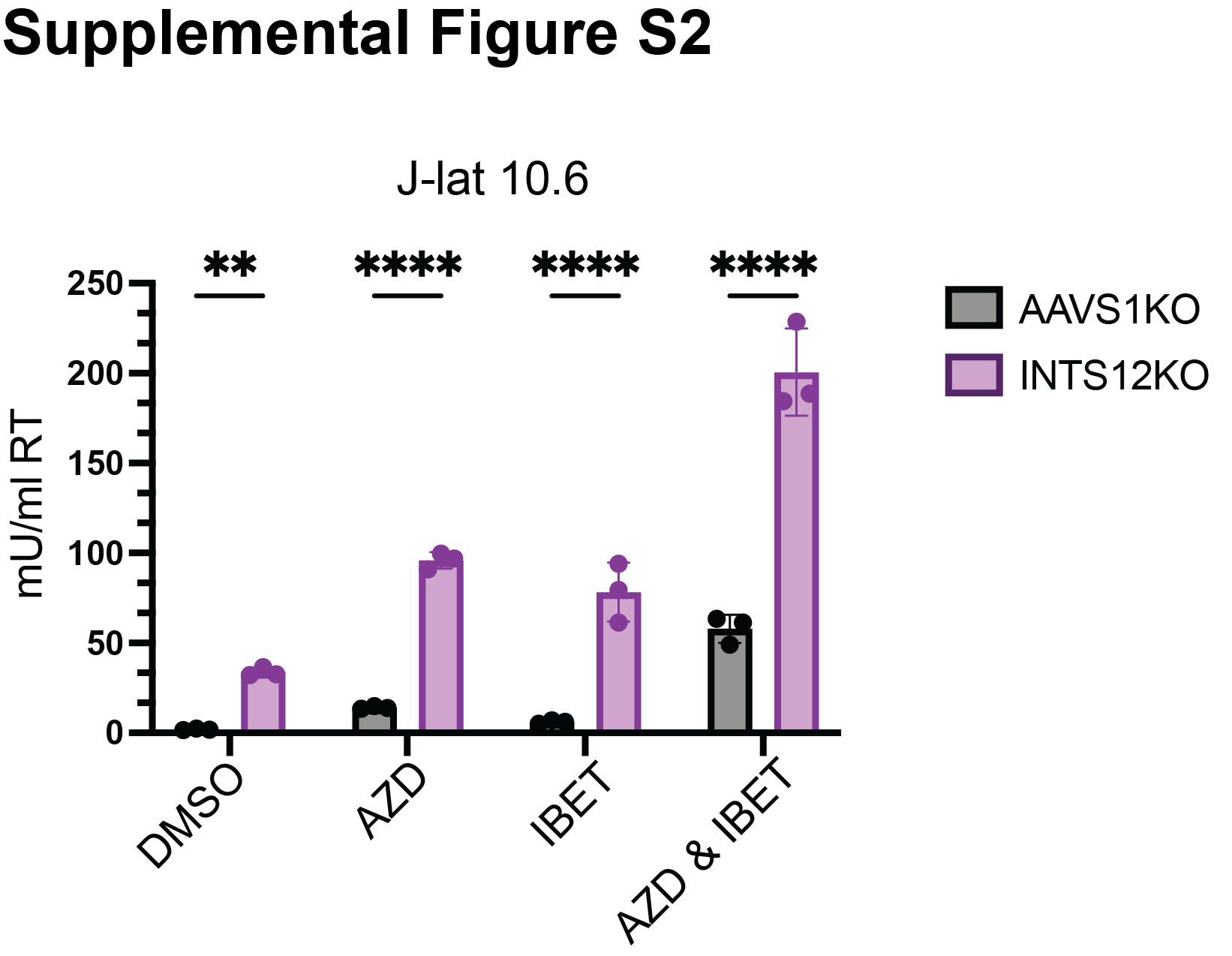

**Figure 2—figure supplement 1**: **Validation of INTS12 knockout in HIV latency reversal both on its own and in the presence of AZD5582 & I-BET151.** J-Lat 10.6 cells were knocked out for INTS12 using the HIV-CRISPR vector with 1 guide targeting INTS12 (cells generated at the same time as Figure 2A). Cells were then treated with 10 nM AZD5582 and or 0.1 µM I-BET151 for 48 hours (or an equivalent volume of DMSO), and HIV reverse transcriptase activity was measured from the supernatant (reported in mU/mL). Untreated = DMSO, AZD = AZD5582, IBET = I-BET151. For statistical analysis, all conditions are compared to the AAVS1 control. 2-way ANOVA, uncorrected Fisher’s LSD, p-value = <0.05 = *, = <0.01 = **, = <0.001 = ***, = <0.0001 = ****.

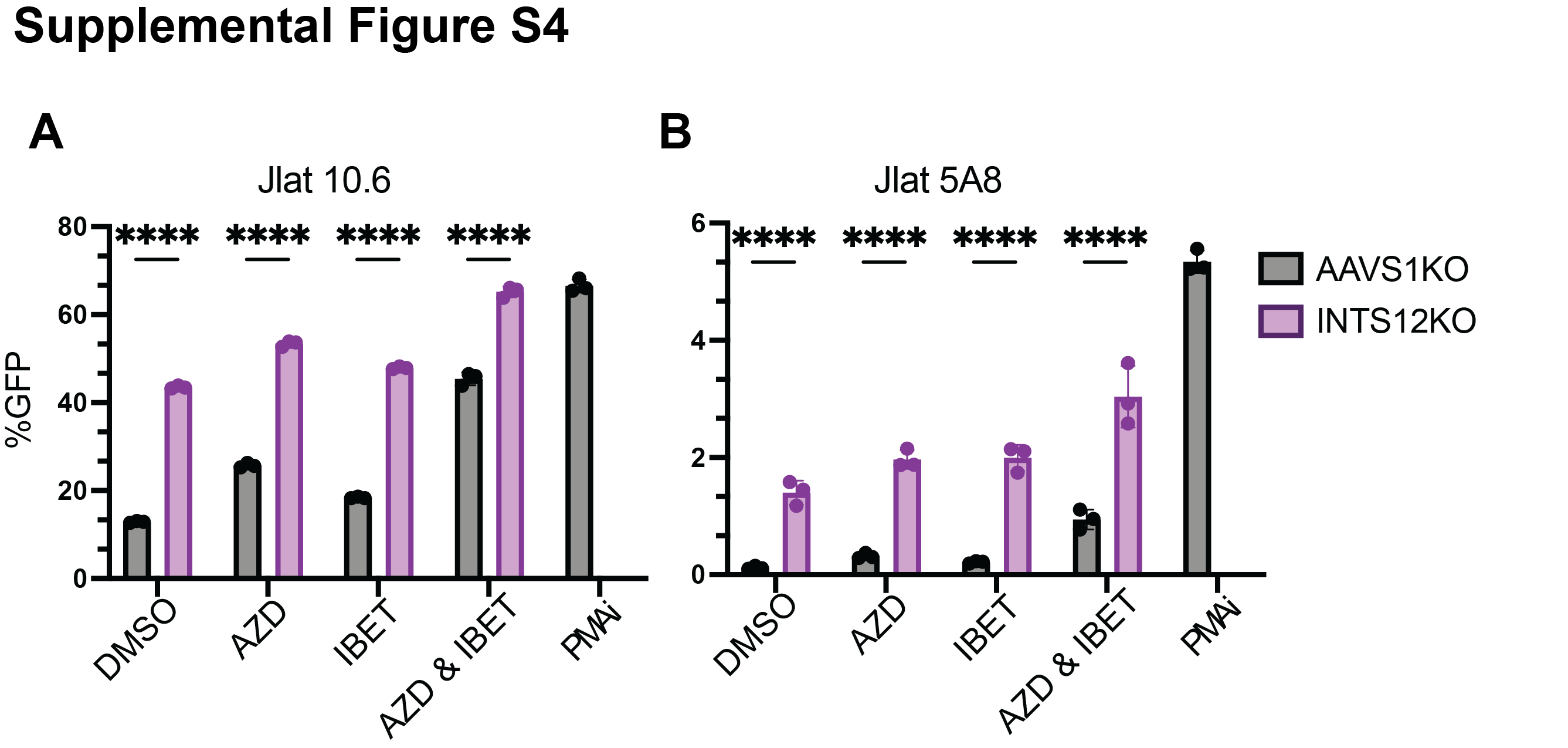

**Figure 2—figure supplement 2**: **Validation of INTS12 knockout in HIV latency reversal both on its own and in the presence of AZD5582 & I-BET151.** J-Lat 5A8 **(A)** and J-Lat 10.6 **(B)** cells were knocked out for INTS12 using the HIV-CRISPR vector with 1 guide targeting INTS12. Cells were then treated with 10 nM AZD5582 and or 0.1 µM I-BET151 for 48 hours (or an equivalent volume of DMSO), and flow cytometry was done to measure %GFP. Untreated = DMSO, AZD = AZD5582, IBET = I-BET151. For statistical analysis, all conditions are compared to the AAVS1 control. 2-way ANOVA, uncorrected Fisher’s LSD, p-value = <0.05 = *, = <0.01 = **, = <0.001 = ***, = <0.0001 = ****.

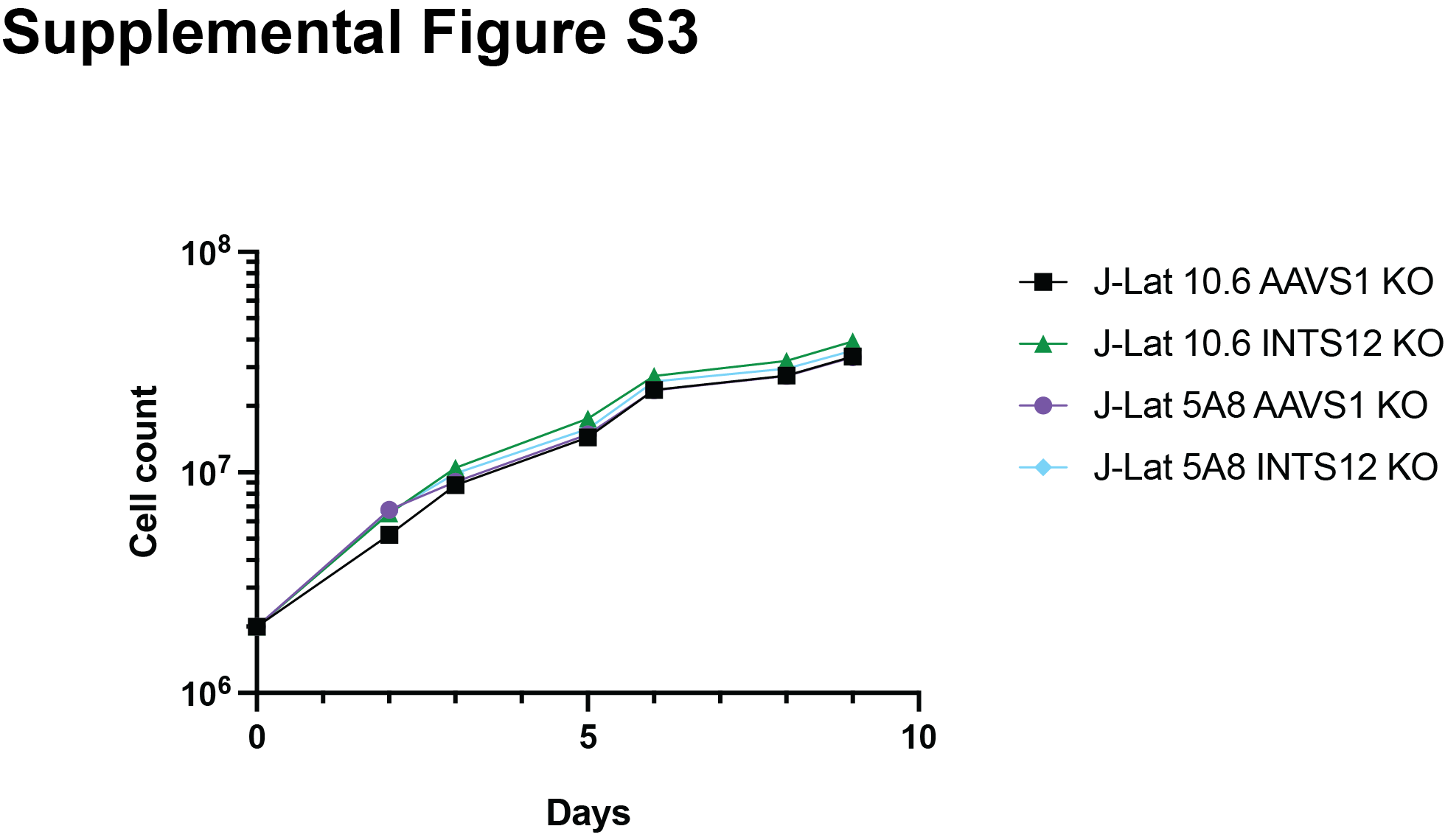

**Figure 2—figure supplement 3**: **Growth Curve of INTS12 knockout cells.** J-Lat 10.6 cells were knocked out for INTS12 using the HIV-CRISPR vector with 1 guide targeting INTS12 or AAVS1 (same cells as Figures 3 and 4). Cells were split every 3 days and counted.

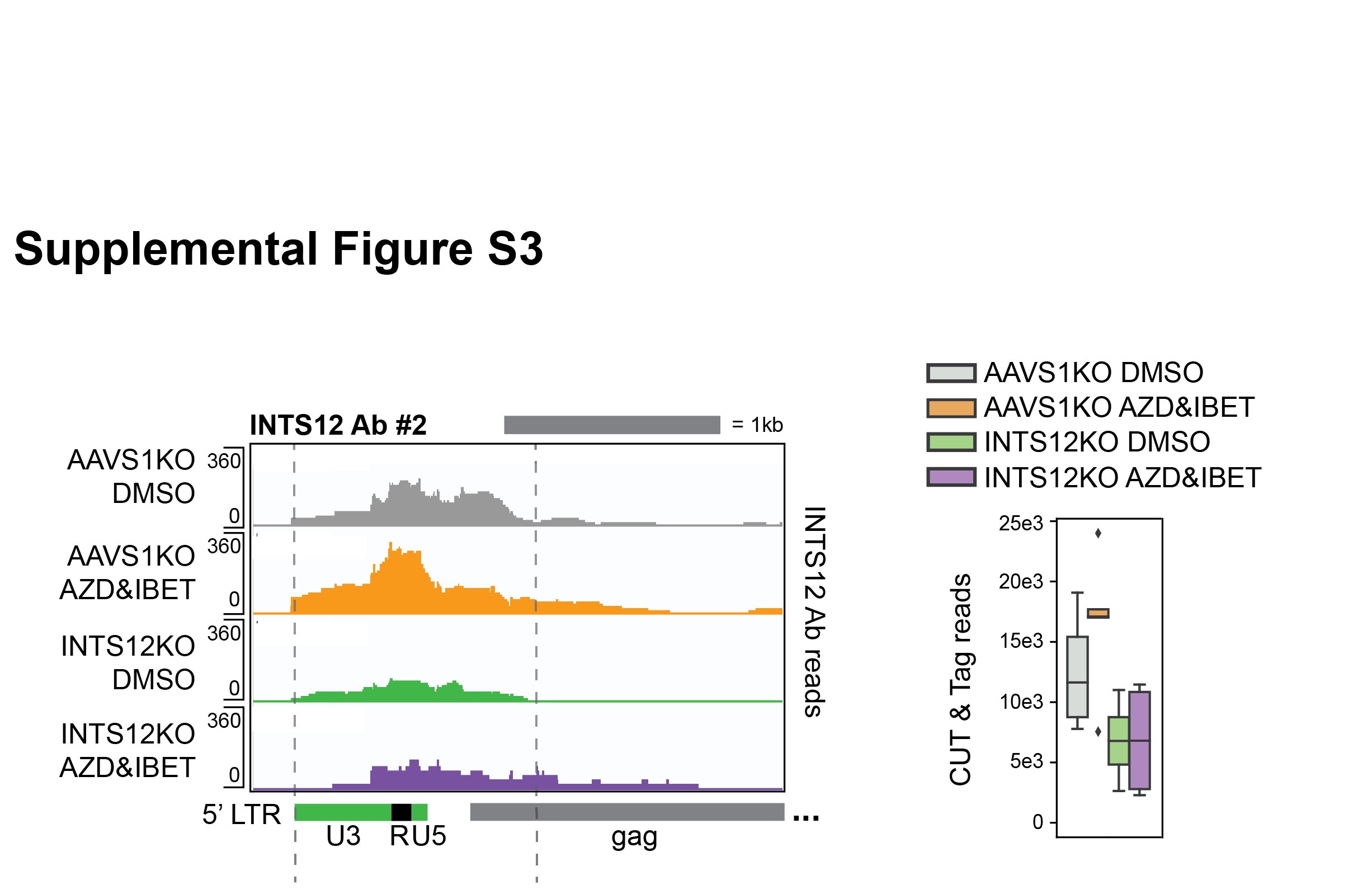

**Figure 5—figure supplement 1**: **INTS12 localization with a second antibody.** Left panel represents pile up graphs of sequencing reads corresponding to INTS12 binding corresponding to the genome location specified below. Right panel is the quantification of all reads between the grey dotted lines on the left. This region matches the same region quantified in Figure 5A for INTS12 Ab #1. All chromosome locations and quantified regions can be found in (supplemental file S4).

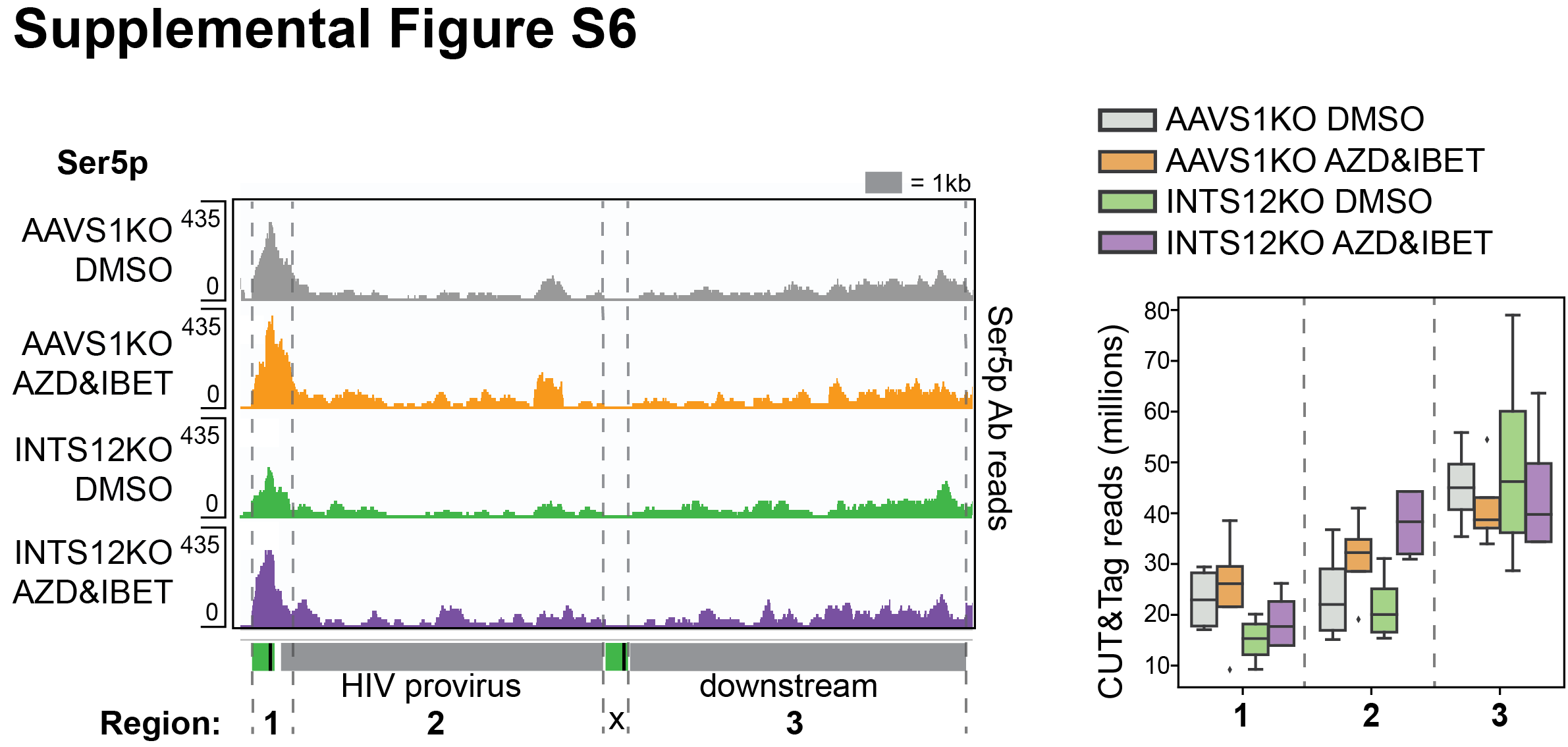

**Figure 5—figure supplement 2**: **Ser5 phosphorylation localization.** Left panel represents pile up graphs of sequencing reads corresponding to Ser5 phosphorylation corresponding to the genome location specified below. Right panel is the quantification of all reads between the grey dotted lines on the left. All chromosome locations and quantified regions can be found in (supplemental file S4).
